# 3D-printing Enabled Micro-assembly of Microfluidic Electroporation System for 3D Tissue Engineering

**DOI:** 10.1101/519496

**Authors:** Qingfu Zhu, Megan Hamilton, Mei He

**Affiliations:** Department of Chemical and Petroleum Engineering, Bioengineering Program, University of Kansas, Lawrence, Kansas, USA; Department of Chemistry, University of Kansas, Lawrence, Kansas, USA

**Author notes:** Electronic Supplementary Information (ESI) is available for multi-directional frequency scanning video*.

## Abstract

Electro-transfection is an essential workhorse tool for regulating cellular responses and engineering cellular materials in tissue engineering. However, existing approaches, including microfluidic platforms and bench top methods, are only able to study monolayer cell suspensions *in vitro*, and are incapable of clinical translation within *in vivo* tissue microenvironment. Knowledge regarding the three-dimensional (3D) electric field distribution and mass transport in a tissue microenvironment is lacking. However, building a 3D electro-transfection system that is compatible with 3D cell culture for mimicking *in vivo* tissue microenvironment is challenging, due to the substantial difficulties in control of 3D electric field distribution as well as the cellular growth. To address such challenges, we introduce a novel 3D micro-assembling strategy assisted by 3D printing, which enables the molding of 3D microstructures as LEGO^®^ parts from 3D-printed molds. The molded PDMS LEGO^®^ bricks are then assembled into a 3D-cell culture chamber interconnected with vertical and horizontal perfusion microchannels as a 3D channel network. Such 3D perfusion microchannel network is unattainable by direct 3D printing or other microfabrication approaches, which can facilitate the high-efficient exchange of nutrition and waste for 3D cell growth. Four flat electrodes are mounted into the 3D culture chamber via a 3D-printed holder and controlled by a programmable power sequencer for multi-directional electric frequency scanning (3D μ-electro-transfection). This multi-directional scanning not only can create transient pores all over the cell membrane, but also can generate local oscillation for enhancing mass transport and improving cell transfection efficiency. As a proof-of-concept, we electro-delivered pAcGFP1-C1 vector to 3D cultured HeLa cells within peptide hydrogel scaffolding. The expressed GFP level from transfected HeLa cells reflects the transfection efficiency. We found two key parameters including electric field strength and plasmid concentration playing more important roles than manipulating pulse duration and duty cycles. The results showed an effective transfection efficiency of ~15% with ~85% cell viability, which is a 3-fold increase compared to the conventional benchtop 3D cell transfection. This 3D μ-electrotransfection system was further used for genetically editing 3D-cultured Hek-293 cells via direct delivery of CRISPR/Cas9 plasmid which showed successful transfection with GFP expressed in the cytoplasm as the reporter. The 3D-printing enabled micro-assembly allows facile creation of novel 3D culture system for electro-transfection which can be employed for versatile gene delivery and cellular engineering, as well as building *in-vivo* like tissue models for fundamentally studying cellular regulatory mechanisms.

## Introduction

Intracellular delivery of regulatory or therapeutic targets into the cell is crucial for pharmacology study as well as the tissue engineering and regenerative medicine.^1–2^ Among various delivery approaches, electro-transfection, also termed electroporation, creates transient permeability of the plasma membrane with temporary pores, due to the external electric field induced high local transmembrane potential. Such an approach has gained increasing popularity due to its safe (chemical free) and effective transfection, and no restrictions on cell types.^3–5^

However, existing electro-transfection systems, including microfluidic platforms and commercial benchtop systems, are only able to study monolayer cell suspensions *in vitro*, and are incapable of clinical translation within *in vivo* tissue microenvironment^6–13^. It has been well documented that cells growing in two-dimensional (2D) culture system significantly differ from living three-dimensional (3D) tissues in terms of cell morphology, functions, cell-to-cell communications, and cell-to-matrix adhesions.^14–15^ Therefore, it is critical to use 3D cultured cells to represent an *in-vivo* like tissue microenvironment. Currently, there is no available 3D culture system to study 3D transfection of tissues. Knowledge regarding the 3D electric field distribution and mass transport in a tissue microenvironment is lacking. Electroporation performed on cell suspensions are very often of limited use when relating to cells within a tissue environment because of the significant variations in terms of membrane compositions, surrounding medium, extracellular matrix, the orientation of cells to the electric fields and so on.^16–17^ Thus, the clinical *in vivo* gene delivery faces tremendous problems.^3, 18^ Although the *in vitro* cellular spheroid model is often applied to study the electro-transfection in a 3D context, these studies only focus on the isolated spheroids or single spheroid and fail to accurately mimic the interactions between cells and the extracellular matrix.^19–20^ Additionally, the mobility of delivered molecules in free solution significantly differs from that in the cellular matrix, and its migration becomes even more difficult when traveling into the cell spheroid.^21^ The investigation of electroporation on 3D cultured cells and tissues has not been explored in the microfluidic platform yet. Benchtop transfection system can only handle single spheroid 3D cells. Herein, we introduce a novel 3D microfluidic electrotransfection system (3D μ-electrotransfection) which allows multidirectional electric field scanning for uniform control of 3D electric distribution, as well as effective nutrient supply for 3D cell culture via the 3D perfusion microchannel network. The 3D μ-electrotransfection system is simply fabricated by the 3D printing-assisted 3D molding and micro-assembling strategy, which employs the LEGO^®^ concept to assemble complicated 3D microchannel networks, facilitating high-efficient exchange of nutrition and waste for 3D cell growth. Such 3D perfusion microchannel network is unattainable by direct 3D printing or other microfabrication approaches. The multi-directional electric field scanning was performed by four flat electrodes mounted into the 3D culture chamber via a 3D-printed holder and a programmable power sequencer. This multi-directional scanning not only can create transient pores all over the cell membrane, but also can generate local oscillation for enhancing mass transport and improving cell transfection efficiency.

As a proof-of-concept, we electro-delivered the pAcGFP1-C1 Vector to 3D cultured HeLa cells in the peptide hydrogel scaffolding for expressing GFP. The critical parameters were optimized including electric field strength, plasmid concentration, pulse duration, and duty cycles. The 3D μ-electrotransfection system was further employed to genetically edit 3D cultured Hek-293 cells via delivery of CRISPR/Cas9 plasmid, which demonstrates the capability and holds the potential for future gene-editing based tissue repair, regenerative medicine, and gene therapy.

## Experimental

### 3D printing and microfabrication of 3D μ-electrotransfection

3D structures were designed and drawn by SOLIDWORKS 2017. The resin mold containing micro-structures were printed by a laptop-sized 3D printer (D3 ProJet 1200, 30-μm resolution) using VisiJet^®^FTX Clear resin (3D systems) for polydimethylsiloxane (PDMS) device production. The Clear resin consists of triethylene glycol diacrylate, sobornyl methacrylate, and 2-3% photoinitiator phenylbis (2,4,6-trimethylbenzoyl)-phosphine oxide as described in the product information. The mold printing followed the reported protocols.^22–23^ Freshly printed molds were cleaned using isopropyl alcohol in sonication and followed with 30-min post cure under UV light. A 20-nm thick palladium or gold coating was deposited onto the surface of the 3D-printed mold using a sputter coater (DENTON, DESK II). Prior to molding, the coated molds were conditioned with the surfactant solution (20% tween 20 in 80% isopropanol) to form a dynamic micellar layer on the metal surface to facilitate the peel-off of polymer microstructures. PDMS was prepared using the standard 10:1 (base to curing agent) ratio. The PDMS mixture was degassed before pouring into the 3D-printed molds and then baked in 40 °C for 12 hours. The 3D printed molds are reusable after cleaning and conditioning. The molded PDMS blocks were then assembled and bound as the 3D μ-electrotransfection device (Fig. 1) after the surface activation using a hand-held corona discharge treater (Electro-Technic Product, Chicago, IL). The assembled 3D μ-electrotransfection device was then bound to a glass slide to complete the fabrication.

**Fig. 1.**
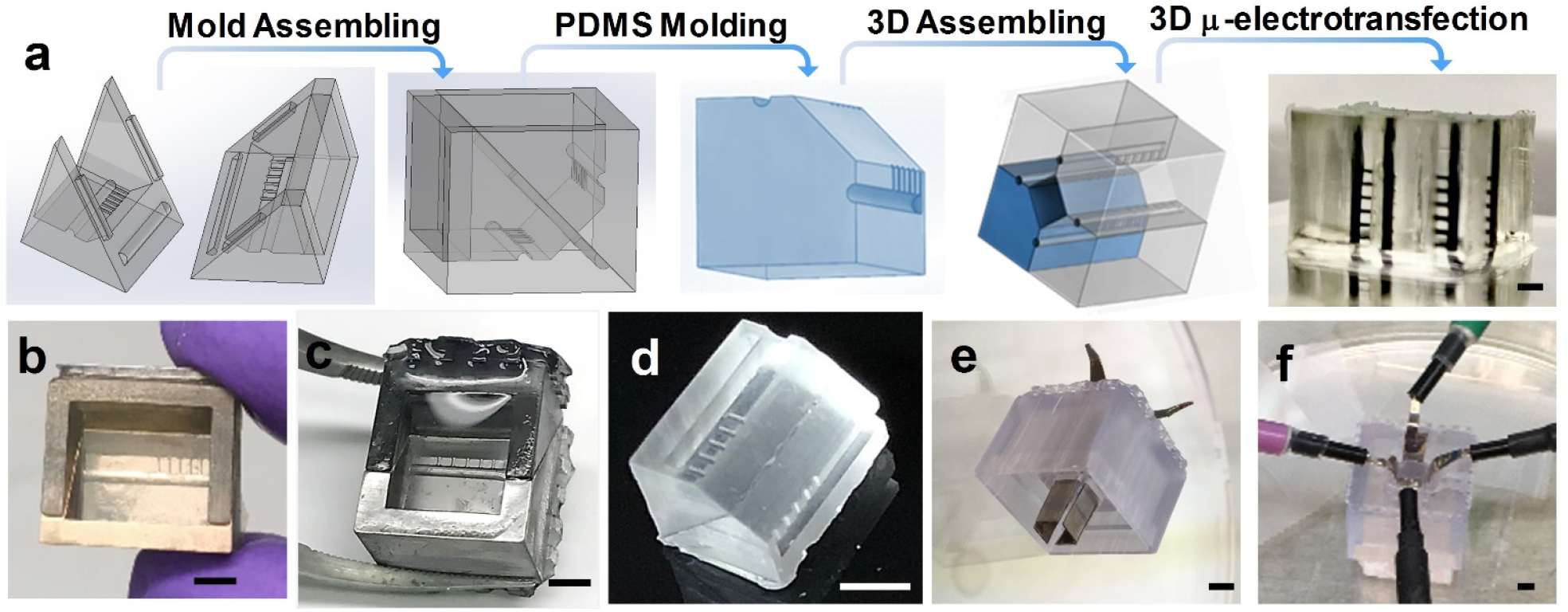
3D printing assisted LEGO^®^ assembling for building 3D μ-electroporation system. a) The concept illustration of 3D printing, molding and LEGO^®^ assembling. Four-piece assembled device bound to a glass slide with channels filled in black dye. The scale bar is 1 mm. b) 3D printed mold (2 pieces assembled) with the surface deposited by 20-nm Au. The scale bar is 2 mm. c) The mold (with 20-nm Ba coating) filled with PDMS and the molded PDMS part is shown in figure d. The scale bar is 2 mm. e) Four electrodes mounted in a 3D printed holder. The scale bar is 2 mm. f) The setup of four electrodes on top of cell culture chip for multi-dimensional electric frequency scanning. The scale bar is 2 mm.

### 3D cell culture and electro-transfection

HeLa cells (ATCC) and Hek-293 cells (ATCC) were cultured and maintained according to ATCC standard protocol with Eagle’s Minimum Essential Medium (EMEM, Sigma-Aldrich), supplemented with 10% (v/v) fetal bovine serum (FBS, Sigma-Aldrich, USA), in a T-25 flask. The peptide hydrogel matrix (PepGel) was used as the scaffold with a peptide gel concentration of 0.2% for 3D cell culture. For culturing 3D cells, the 2D cells growing at a confluency of 80-90% were re-suspended by 0.25% trypsin/EDTA (Sigma-Aldrich, USA) solution and centrifuged for 5 min at 250 g-force for seeding into peptide hydrogel. The seeding density for 3D cell culture was ~2.5 × 10^5^ cells/mL. The 3D cell culture was carried out following the protocol described in the literature.^24^

A 4.7 Kb plasmid pAcGFP1-C1 (Clontch, Mountain View, CA) encoding green fluorescent protein (GFP) was amplified in NEB^®^ 5-alpha Competent E.coli (New England Biolabs, Ipswich, MA) and isolated by QIAGEN Plasmid Maxi kit (QIAGEN GmbH, Germany). The plasmid purity was determined using Nanodrop 2000 spectrophotometer. A 9.2 Kb CRISPR /Cas9 vector pSpCas9(BB)-2A-GFP (PX458) (Addgene, MA, USA) was amplified and purified following the same protocols as the preparation of GFP plasmid. The 2D standard electroporation protocols (Neon^®^ transfection system) were used for validation of 3D μ-electrotransfection system. The successfully transfected cells can express GFP as the reporter and be analyzed under flow cytometry (BD FACSAria IIIu, BD Biosciences, USA) and confocal microscopy (Olympus IX81/3I spinning disk confocal inverted microscope).

### 3D COMSOL simulation

The AC/DC module was applied to simulate electric field distribution in 3D μ-electrotransfection system using COMSOL Multiphysics software package (COMSOL Multiphysics 5.2). The equation governing electrostatics was numerically solved for the device to arrive at steady-state solutions. At steady state, the electric currents in a conductive media is given by the Ohm’s law, which states

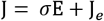

where *σ* is the electrical conductivity, *ϕ* is the electric potential, **J** is the current density and **E** is the electric field. Electric field distribution was visualized using the expression shown below,

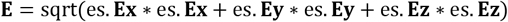

where es.Ex, es.Ey, and ex.Ez are the components of the electric field in the dimension of x, y, z, respectively. The conductivity of electroporation buffer was set at 0.14 S/m.^25^ The relative permittivity of peptide hydrogel was set at 1, based on literature for a similar peptide hydrogel.^26^ We carried out electrostatic numerical simulations to predict the distribution of electric field strength across the entire cell culture chamber. The multi-directional electric frequency scanning was simulated and performed following the protocol as shown in Fig. 2b. To simulate nutrient medium exchange and diffusion in 3D μ-electrotransfection system, the transport of diluted species in porous media model was studied in a time-dependent manner. Three different cases, i.e. medium diffusion from top medium to cell matrix, medium diffusion from side perfusion microchannel network to cell matrix, and medium from both top and side perfusion microchannel network to cell matrix were investigated for comparison.

**Fig. 2.**
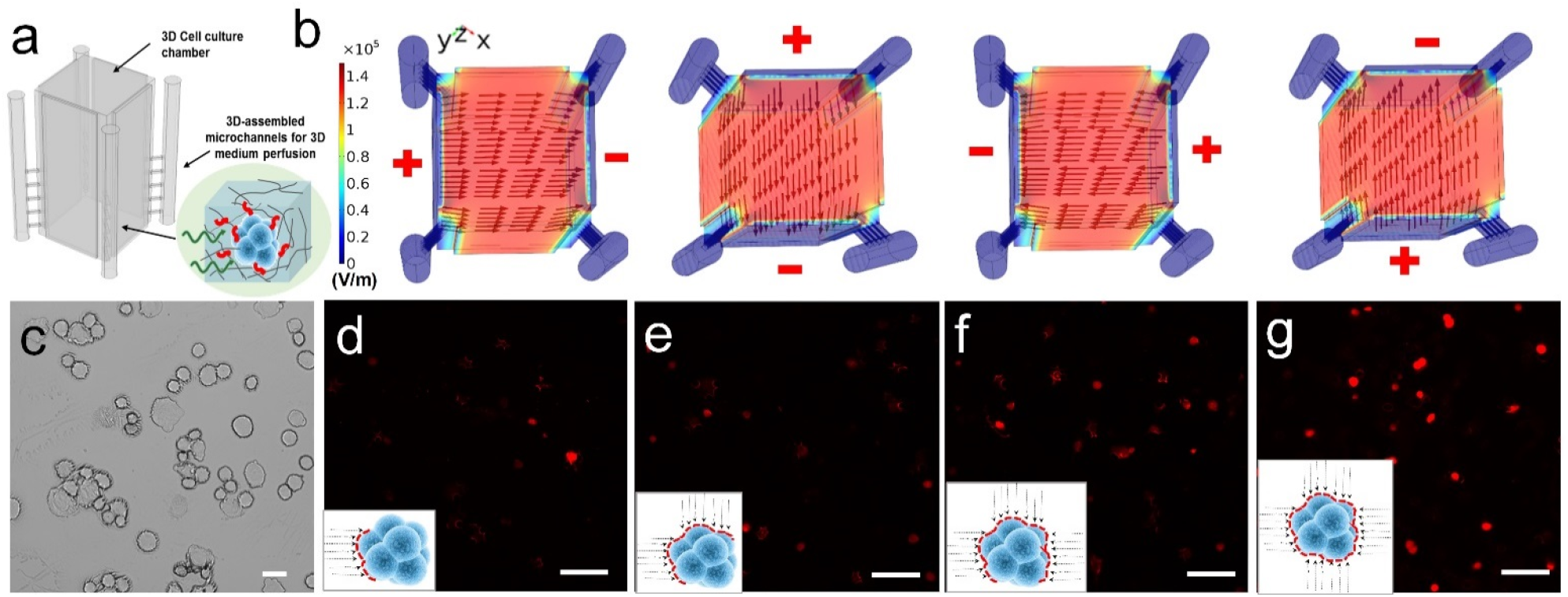
a) Illustration of on-chip 3D cell culture. b) COMSOL simulation of electric field distribution across the cell culture chamber with the multi-directional field scanning strategy. A voltage of 500 V was applied. Arrows indicate the intensity and direction of the electric field. c) Control HeLa cells without applying electric field under bright field. The scale bar is 20 μm. d) one-direction electric filed based PI delivery. e) two-direction electric filed based PI delivery. f) Three-direction electric filed based PI delivery. g) Four-direction electric filed based PI delivery. The scale bar is 50 μm for Fig. 2 d to g.

### 3D multi-directional electro-transfection

The voltage across the 3D cell culture chamber in multi-directional scanning was programmed and delivered by HVS448 800 high voltage sequencer (LabSmith, CA, USA), which generated a square-wave electric pulse (see Supplemental Video). Prior to applying the voltage, the cell medium (EMEM) was removed and 100 μL low conductivity medium (Cytoporation^®^ Medium T, BTX) was added on top as well as in the side perfusion channels. The electric potentials were applied to flat electrodes (Ag/Pa alloy, 3.5 mm width and 0.2 mm thick, Fisher). The electrodes were mounted by a customized holder, which was printed by a 3D printer (D3 ProJet 1200) as shown in Fig. 1e. For transfection, the GFP plasmid was prepared in the gel matrix with cells to achieve a final concentration between 60 to 140 μg/mL. Cell culture was kept on ice during electroporation. Pulses at a frequency of 1Hz were applied with a given electric field intensity. After electroporation, low conductivity medium in cell culture was replaced by EMEM medium (pre-conditioned in 5% CO_2_). The transfected 3D cells were cultured for two days and assessed for GFP expression levels by fluorescence microscopy and flow cytometry.

### Flow cytometry analysis for assessing 3D electro-transfection efficiency and cell viability

The 3D transfected cultures were transferred from chip to a 1.5 mL centrifuge tube according to the method described in the PepGel protocol. Without removing the upper layer cell medium, the gel was mechanically disrupted thoroughly by pipetting up and down. The mixture was centrifuged at 600 g for 6 min. To break colonies, 100 uL accumax^TM^ solution (Sigma-Aldrich, USA) was added and incubated for 5 min at 37 °C. To stain the dead cells, 5 μL of 10 μg/mL propidium iodide (PI, Sigma-Aldrich, USA) was added and incubated in dark for 2 min. Thereafter, the mixture was centrifuged at 200 g for 5 min and the cells were re-suspended with 200 μL PBS containing 0.5% BSA (w/v) for flow cytometry analysis. A total of 5000 events were measured in each sample at a flow rate of approximately 80 events/s.

## Results and Discussions

### 3D-printing enabled micro-assembling of 3D μ-electrotransfection system

To build a 3D microfluidic electro-transfection system capable of uniform 3D electric field distribution as well as acting as an effective 3D cell culture device, we conceived an electroporation chip as shown in Fig. 1. Conventional microfabrication approaches are unable to construct such 3D microstructures, due to complicated protocols for accurate alignment and multilayer bonding. Even though, it is very challenging for direct 3D printing of monolithic 3D microstructures, particularly micro-scale hollow channels.^27–28^ Therefore, we introduced a 3D-printing assisted molding of PDMS as LEGO^®^ blocks for assembling into a more complicated 3D device. As illustrated in Fig. 1a, two designed parts were assembled as one transfer mold for PDMS molding. The assembled mold is detachable for easy release of PDMS parts. The PDMS polymer was completely cured in the mold and no microstructural defects were identified during the demolding process (Fig. 1d). It is worth mentioning that high-temperature baking (e.g. > 40 °C) should be avoided as high temperature may cause physical structural distortion of 3D printed resin. The molded PDMS polymer replicates microstructures as a single assembling unit shown in Fig. 1c and d. After four units are assembled with permanent bonding using surface plasma treatment, the 3D microfluidic electrotransfection device can be formed with four main vertical microchannels (~350 μm) each connected with five horizontal microchannels (~200 μm) (Fig. 1a). To facilitate the precise production of microstructures, sputtering Ba or Au coating was deposited onto the mold surface in 20-nm thick as shown in Fig. 1b and c. The final assembled device can be bound onto glass slide after surface plasma treatment. The electrodes were fixed in a 3D printed holder (Fig. 1e), which fits into the central culture chamber for electroporating 3D cultured cells (Fig. 1f and Fig. 2a).

### Working principles of multi-directional electric field scanning enabled 3D transfection

In an electric field across cells, the cell membrane is an electric insulator that separates extracellular medium with the intracellular medium. The ion concentration gradient between outside and inside of the cellular membrane generates a resting potential difference, which is homogenous all along the cell membrane.^29^ Upon application of voltage, such electric potential difference across the cellular membrane will disrupt the field lines,^30,31^ consequently, leading to the current forced to flow around the cell and forming the ionic layers along the cellular membrane. The largest field line distortion is recognized at the sides of the membrane facing the field lines.^3, 32^ After reaching a certain intensity of field strength, the cell membrane can be disrupted to create transient pores. In a uniform electric field, the induced potential difference (Δ*Ψ*) at a point on the cell membrane and at a time after the rise of the electric pulse is given by^3, 33^:

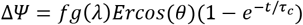

Where *θ* is the angle formed between the direction of the electric field and a normal point on the membrane. *f* is a factor related to the shape of the cell which equals to 1.5, if the cell is spherical. *g(γ*) is a factor related to the conductivity of the membrane. *r* is the semi-axis aligned along the electric field and *τ_c_* is the charging time of the cell membrane. Because the membrane conductivity is extremely low compared to the conductivity of the intra and extracellular medium, it can be assumed that *g(λ*) = 1. As the electric field pulse durations (*τ_c_*) is very small, at a steady state, considering the cell as a spherical insulator shell, ΔΨ can be written in a simplified expression:

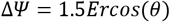

Therefore, the induced potential difference on the cell membrane is directly proportional to the cell size and the strength of the electric field. Furthermore, the resting potential difference across the cell membrane reaches its maximum value at the side of the membrane facing the electric field directly (0° or 180°), and decreases progressively along the cell surface up to the poles. Therefore, to open more pores on cell membrane with mild voltage conditions, changing electric field directions to create pores from multiple sites on the cell membrane is more effective for electroporation. Therefore, in our experiments, the multi-directional electric frequency scanning method was developed, which can easily create transient pores all over the cell membrane along the multidirectional scanning as demonstrated in Fig. 2b, as well as create local oscillation for enhancing mass transport and improving cell transfection efficiency. By COMSOL simulation in Fig 2b, the uniform electric field strength across the cell chamber is estimated at 1000 V/cm with a voltage of 500 V, which is sufficient to create transient pores on the cell membrane. In general, the electric field across cell clusters needs to reach the electroporation threshold (100 to 1000 V/cm, depending on cell type) allowing membrane disruption.^3^

To demonstrate the effectiveness of multi-directional electric field induced membrane disruption for delivering targets into the cytoplasm, a small molecule dye PI was electro-delivered to investigate the electropermeabilization. PI cannot permeate living cells with an intact cell membrane, but can enter into cytoplasm once the cell membrane is disrupted. As demonstrated in Fig. 2d to g, the electric field has been manipulated from one to four directions while maintaining constant electroporation duration. The fluorescence of cytospined cells on the glass slide was tracked by the microscopy analysis to evaluate PI delivery efficiency. A transfection voltage of 400 V with four electric pulses (16 ms) at a frequency of 1 Hz was applied. The single direction of electric field only gives ~ 40% cells transfected, which is much less than four directional scanning with over 80% cells transfected.

### 3D μ-electrotransfection system for 3D cell culture

Unlike 3D cell cultures in the well plate where medium exchange only takes place from the top of well plates, our system allows multidimensional diffusion of the fresh medium from both the top of cell matrix and the side of microchannel arrays surrounded the culture chamber in vertical and horizontal directions. Such 3D perfusion microchannel network allows better nutrient support and waste exchange needed for effective growth of 3D cells and tissues. The mass transport and efficient diffusion of nutrients from multidirections were demonstrated by COMSOL simulation in Fig. 3, which indicates that a 3D perfusion microchannel network significantly improves the medium exchange compared to the conventional 3D cell culture in the well plate.

**Fig. 3.**
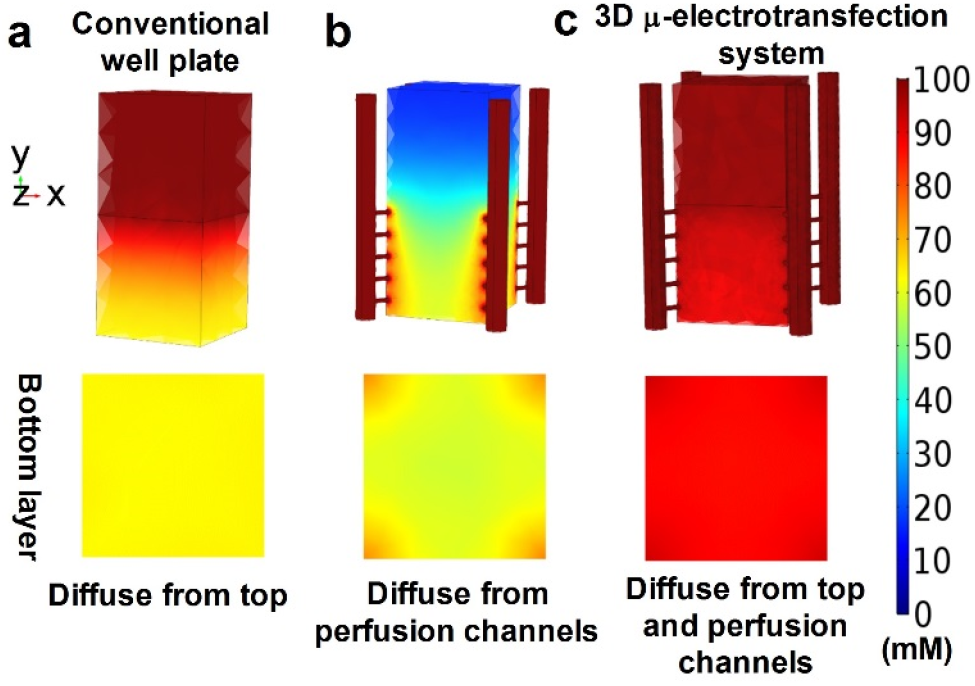
COMSOL simulations of medium diffusion. a) Diffusion from top medium to cell matrix in the vertical direction for the conventional well plate. b) Diffusion from microchannel network to cell matrix in the horizontal directions. c) Diffusion from both microchannel network and top medium. 100 mM ionic strength was used in the fresh medium for simulation.

We chose the peptide hydrogel as scaffolding for 3D cell growth in our 3D μ-electrotransfection system shown in Fig. 4a, due to its high encapsulation stability, cell attachability and biocompatibility.^34–35^ Upon crosslinking, the hydrogel forms a porous matrix with a pore size ranging from 200 to 400 nm, which gives a stable physical support for 3D cell growth as imaged in Fig. 4 e and f.^24^ Due to the perfusion microchannels for nutrients and waste exchange in our device, a high cell seeding density from 1×10^5^ to 5×10^5^ cells/mL can be achieved. The cell showed excellent attachability to peptide fibers (Fig. 4f). The dense spheroid distribution in peptide hydrogel matrix in 3D as well as the morphology of a single spheroid have been characterized by confocal microscopy shown in Fig. 4 b to d. An individual 3D spheroid with the size of ~ 50 μm was observed after culturing for 4-5 days which is composed of ~ 10 cells as shown in Fig. 4c and 4d. Such morphology characterization indicates that our 3D μ-electrotransfection system provides a suitable microenvironment for growing 3D cells and tissues, which is enabled by the implementation of the 3D perfusion microchannel network.

**Fig. 4.**
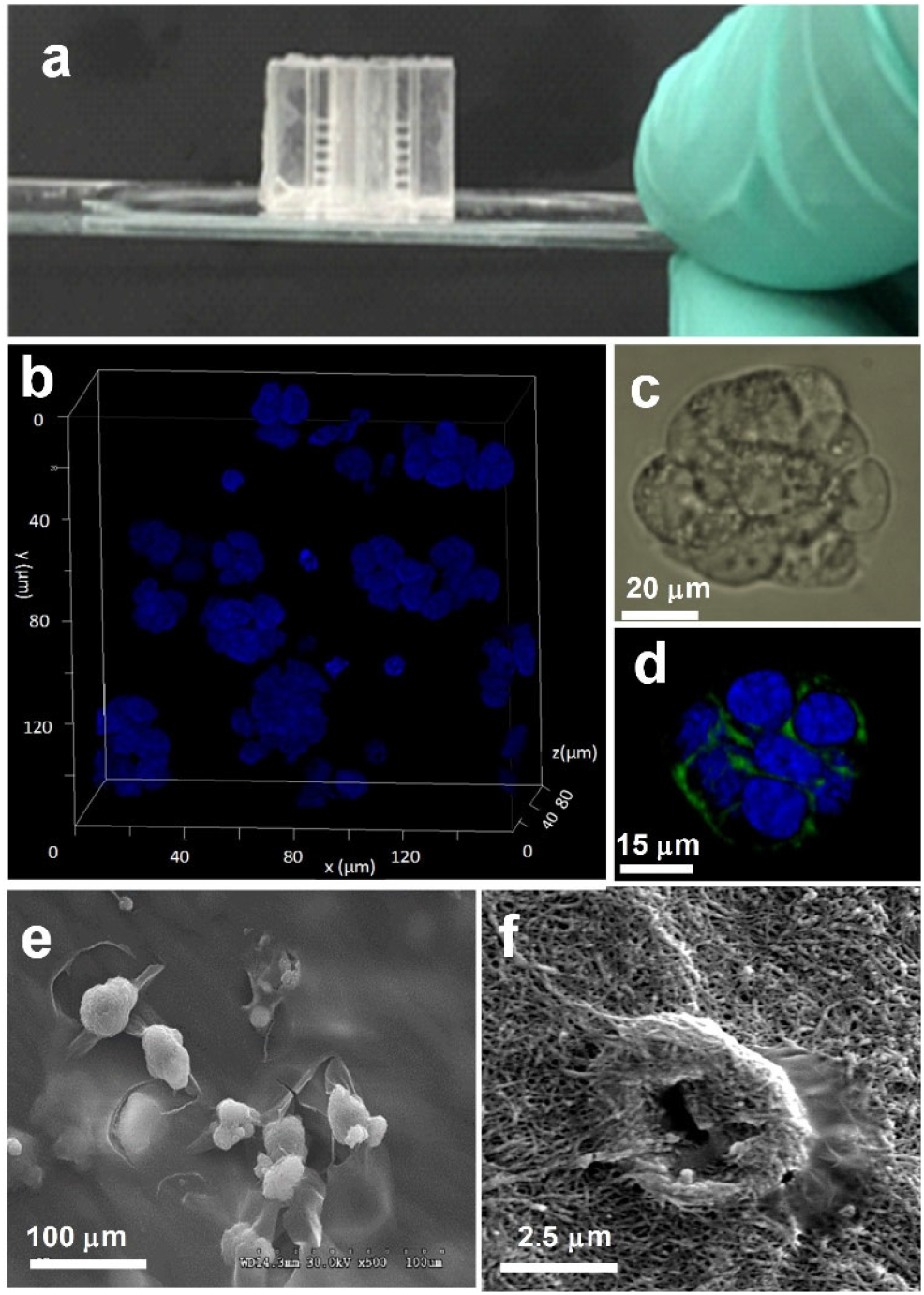
On-chip 3D cell culture and microscopy characterization. a) Assembled PDMS chip bound to a glass slide for cell culture. b) Distribution of 3D-cultured HeLa cell spheroids in peptide hydrogel under confocal microscope imaging. Nuclear DNA was stained by Hoechst. c) The close look of a single 3D cell spheroid in bright field and in the fluorescence channels in d. The cell membrane was stained by FITC conjugated antibody-dye and the nuclear DNA was stained by Hoechst. e) SEM imaging the cell spheroid distribution and the single spheroid interacting with peptide fibers in f.

### 3D μ-electrotransfection of 3D cultured cells

To assess the transfection efficiency and cell viability of our 3D μ-electrotransfection system, a plasmid DNA (pAcGFP1-C1) encoding GFP was electroporated into 3D cultured HeLa cells. For conventional electrotransfection of 2D cell suspension, the critical electric field needs to reach a value in the range of 100 to 1000 V/cm (depending on cell size and electric field property) to disrupt the cell membrane and ensure the reversibility.^3, 36^ In the case of spheroid cells, the low electric field (~500 V/cm) with long pulses (~20 ms) has been reported to lead to transfected GFP expression.^19, 37^ In contrast to the study of cell suspensions or isolated cell spheroids, we intended to deliver plasmid directly to 3D cells embedded in the extracellular matrix to mimic electroporation of *in vivo* like tissue microenvironment. Thus, we optimized key parameters that control the electroporation efficiency, including electric field strength, plasmid concentration, pulse duration, and duty cycles.

With a voltage of 300 V in multi-directional field scanning (equal to an electric field of 750 V/cm), 2.5% GFP expressed cells were identified from the total cell population with cell viability of ~ 95% by flow cytometry analysis (Fig. 5). The transfection efficiency was increased with an increase in the applied voltage from 300 V to 600 V. However, cell viability decreased from ~ 96% to ~ 84%, due to higher more dead cells caused by the high voltage, which in turn decreases the transfection efficiency with a higher voltage of 600 V (1500 V/cm) (Fig. 5 a). Either increasing the pulse duration or adding more duty cycles can lead to the increase of higher cell death rates caused by high voltage, but the cell death rate dramatically increased accordingly, due to the harsh, irreversible electric interruption of cell membrane. The 4 ms pulse duration resulted in transfection efficiency of 15.6%, with 58.3% cell viability. Increasing the plasmid concentration will create more contact opportunities between cells and plasmids, reflecting an increased number of transfected cells. The optimal plasmid concentration appears to be ~120 μg/mL, after which increasing the plasmid DNA concentration did not result in a higher transfection efficiency. This observation is in agreement with a previous report that there is a maximum plasmid concentration for gene delivery.^38^ Compared to 2D cell transfection, the optimal plasmid concentration for 3D cell electroporation is much higher (110 vs 40 μg/mL).^39^ This is attributed to the porous peptide hydrogel matrix which limits the travel of plasmid to cells and requires more amount of plasmid to enhance contact opportunities with cells. To find the balance between the transfection efficiency and cell viability for the best transfection outcome, the optimized voltage is 500 V with ~120 μg/mL plasmid concentration, parameters which were shown to play a more important role in the control of good cell viability, compared to the electric duty cycle and pulse. In addition to the scaffold matrix effect, the 3D cell spheroid is much bigger than an individual cell and the plasmid needs to travel a long distance to reach the cells inside the cluster, which makes the gene delivery even more difficult. Compared to the reported data,^40^ our method showed ~ 3 fold higher transfection efficiency with high cell viability (> 85%), which is attributed to the electric filed scanning strategy. Fig. 5b to e shows a typical image and flow cytometry analysis of 3D cultured cells after electro-transfection in our system. The confocal microscopic imaging in Fig. 5d displays a transfected 3D cell spheroid with a diameter around 60 μm.

**Fig. 5.**
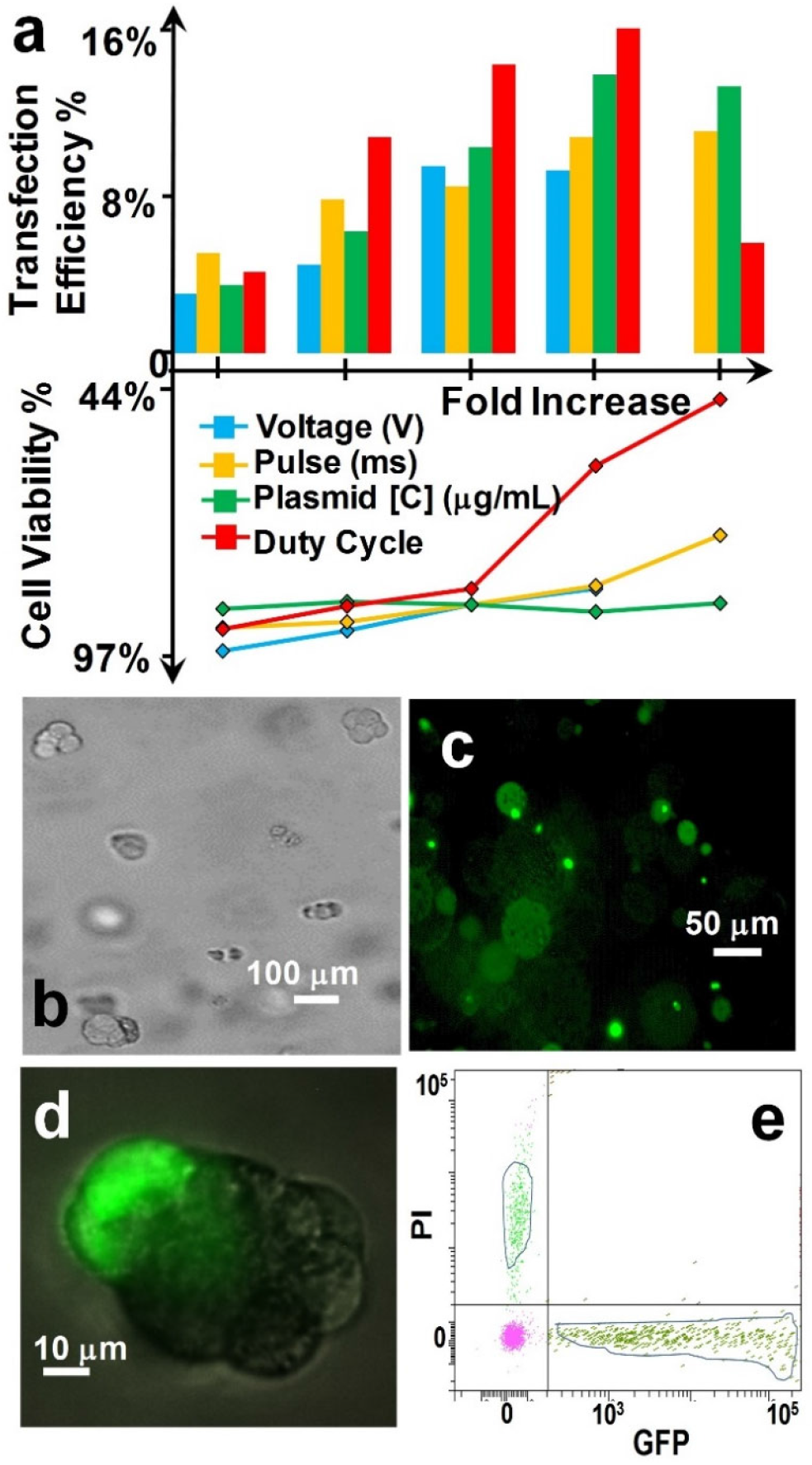
Electro delivery of GFP plasmid to 3D cultured HeLa cell spheroid within peptide hydrogel. a) Investigation of key parameters including voltages (300 400, 500, 600 V), pulse duration (0.4, 1, 2, 3, 4 ms), plasmid concentration (60, 80, 100, 120, 140 μg/mL) and duty cycles (1, 2, 3, 4, 5) on transfection efficiency and cell viability. b) Electroporated sample under the bright field and FITC channel in c. The transfection voltage was 500 V with a pulses duration of 3 ms for each direction, at the frequency of 1 Hz, and 3 duty cycles. d) Confocal image of a transfected cell spheroid. e) Evaluation of GFP positive cell and PI positive cell via flow cytometry.

For proving the applicability of our system in 3D tissue engineering, we transfected the CRISPR/Cas9 gene with 3D cultured Hek-293 cells. The CRISPR/Cas9 editing has emerged as a rapid and powerful approach to make precise and targeted changes to the genome of living cells.^41^ Recent studies have successfully demonstrated gene transfer to organoids by various methods including lentivirus transfection,^42^ liposomal transfection^43^ and electroporation.^44^ The advantage of electroporation over other methods is no need of chemicals required in the culture system. The plasmid can be delivered upon preparation and does not require production of lentivirus or carriers, which significantly reduces the labor and time. In this experiment, a 9.2 Kb PX458 vector was delivered to 3D cultured HeK-293 cells using our 3D μ-electrotransfection system. The EGFP protein encoded by this CRISPR/Cas9 vector was traced to evaluate the CRISPR/Cas9 delivery. The high concentration of CRISPR/Cas9 plasmid (200 μg/mL) was applied according to the previously optimized conditions, and a mild voltage (400 V)) and 2 duty cycles were chosen. Fig. 6a shows the 3D distribution of transfected Hek cells within the cultured extracellular matrix. Fig. 6b shows the successfully transfected cell spheroids with ~10 cells in a perfect round cluster. Fig. 6c shows a single transfected cell in the status of the division. The uniform green fluorescence distribution within either the spheroids cluster or the divided two daughter cells illustrates the effective electroporation across overall cell cytoplasm.

**Fig. 6.**
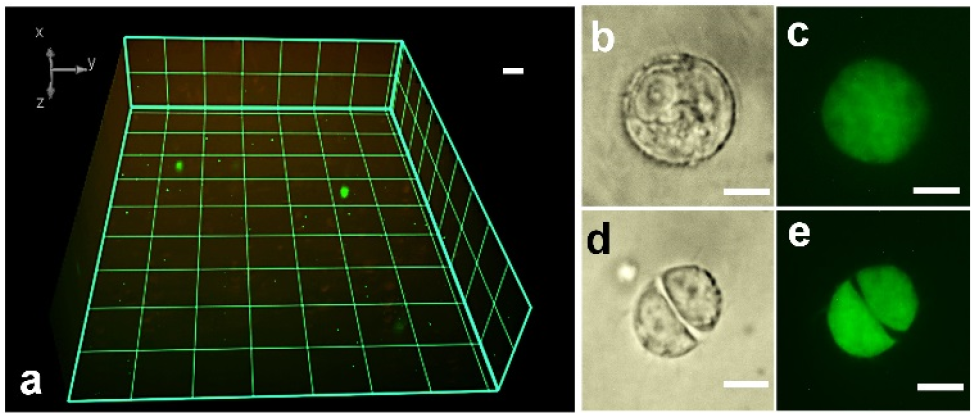
Electroporating the CRISPR/Cas9 plasmid (PX458) to 3D cultured Hek-293 cells. a) The overview of the transfected cells in the 3D matrix. b) A transfected Hek-293 cell spheroid under the bright field and in FITC channel in c. The scale bar is 20 μm. d) A single transfected cell in the process of the division under the bright field and the FITC channel in e. The scale bar is 10 μm.

## Conclusions

In recent years, 3D printing has drawn much attention from the research community, due to the capability of creating complex structures with high quality for fast prototyping^45–47^, compared to traditional micro-fabrication. As a layer-by-layer manufacturing method, 3D printing not only achieves monolithic device fabrication but also allows for printing molds used in producing a PDMS microfluidic chip. However, constructing complex 3D structures and monolithic hollow channels in micro-scale is still challenging. In this paper, we take advantages of 3D printing and a LEGO^®^ assembling concept to construct more complicated 3D microfluidic channels, which extends the 3D printing capability for creating unattainable 3D micro-geometries and introducing geometrics enabled functionality. Such assembled 3D μ-electrotransfection system allows spatial and temporal control of electric field uniformly in three dimensions. Therefore, the multi-directional electric frequency scanning is achievable for maximizing the electroporation efficiency via enhancing the resting potential difference across all over the cell membrane. Furthermore, this scanning process also creates local oscillation for enhancing mass transport and improving cell transfection efficiency.^48^ The 3D-cell culture performance is improved as well due to the enhanced medium perfusion via interconnected vertical and horizontal perfusion microchannels array, which is reconstructed by this 3D printing-assisted molding and assembling process.

Existing microfluidic electroporation approaches are only able to study monolayer cell suspensions *in vitro*, which is incapable of clinical translation within *in vivo* tissue microenvironment but essential in gene therapy and tissue repair. This is the first 3D microfluidic electroporation system for transfecting 3D cultured cells, which demonstrated a ~3-fold increase of transfection efficiency with good cell viability (> 85%) compared to the conventional benchtop 3D cell transfection. The optimization of several key parameters, including electric field strength, plasmid concentration, pulse duration and duty cycles, gave a good rationale for understanding the influence on the delivery process and cell viability. The threshold of permeabilization voltage and plasmid concentration play more important roles, due to the direct connection with the chances of transient pore opening and contact. This study also can mimic the intracellular delivery of therapeutic molecules *in vivo* and has important implications for gene delivery in tissues, especially for editing cells *in vivo* using the CRISPR/Cas9 method. Future work will be conducted to further enhance the transfection efficiency by investigating different scaffolds with various conductivity and porosity. This 3D re-constructed μ-electrotransfection platform can serve as a model system for studying and mimicking *in vivo* electro-transfection process, and building the foundation for developing more effective clinical gene delivery approaches.

## Acknowledgments

Dr. Qingfu Zhu performed overall experiment, data collection and analysis, as well as drafted the manuscript. Miss Megan Hamilton performed 3D cell culture, plasmid amplification, and transfection. Dr. Mei He conceived the research idea, analyzed the data, and revised the manuscript. We acknowledge the funding support from USDA-NIFA KS451214 and NIH P20 NIH0078730, as well as the K-INBRE Postdoctoral Scholarship (P20 GM103418) for QZ. We also thank the assistant from the Confocal Core at the University of Kansas and the Kansas State University for imaging 3D cells.

